# Emergence of variation between groups through time in fish shoal collective motion

**DOI:** 10.1101/2021.02.23.432454

**Authors:** Hannah E. A. MacGregor, Christos C. Ioannou

## Abstract

Despite extensive interest in the dynamic interactions between individuals that drive collective motion in animal groups, the dynamics of collective motion over longer time frames are understudied. Using three-spined sticklebacks, *Gasterosteus aculateus*, randomly assigned to twelve shoals of eight fish, we tested how six key traits of collective motion changed over shorter (within trials) and longer (between days) timescales under controlled laboratory conditions. Over both timescales, groups became less social with reduced cohesion, polarisation, group speed and information transfer. There was consistent inter-group variation (i.e. collective personality variation) for all collective motion parameters, but groups also differed in how their collective motion changed over days in their cohesion, polarisation, group speed and information transfer. This magnified differences between groups, suggesting that over time the ‘typical’ collective motion cannot be easily characterised. The minimal sample size of independent groups and their divergence over time need to be considered in future studies.

## Introduction

Collective motion is widespread in living systems from bacteria to humans (Deutsch et al., 2012; Vicsek and Zafeiris, 2012). In these systems, there is no centralised control of group movement yet highly synchronised behaviours can arise from local interactions between individuals, even when individuals lack an awareness of the position and movement of all group members (Giardina, 2008). Parallels are often drawn between animal collective behaviour and physical and chemical systems during phase transitions (Nicolis and Prigogine, 1977; Vicsek et al., 1995), however the traits of animal groups (e.g. their speed, direction, and spatial organisation) differ fundamentally from non-living systems in that they emerge from interactions between complex and biologically-motivated individuals and are shaped by natural selection (Sumpter, 2006). Although there are costs to animal collective behaviour as it can result in competition for food (MacGregor et al., 2020) and exploitation by predators and parasites (Bauer et al., 2015; Rode et al., 2013), the independent evolution and diversity of collective behaviours across animal taxa suggest that benefits from improved predator avoidance (Ioannou et al., 2019) and use of resources (Sasaki et al., 2013) often outweigh the costs.

Shifts in the benefits and costs of collective behaviour can result in changes over evolutionary time (Greenwood et al., 2013; Kowalko et al., 2013; Wood and Ackland, 2007). Over within-generation timescales, collective behaviour can change in response to changing environmental conditions, such as cues from predators or food, or abiotic factors such as habitat structure (Chamberlain and Ioannou, 2019; Hoare et al., 2004; Ling et al., 2019; Rodriguez-Pinto et al., 2020; Romenskyy et al., 2020; Schaerf et al., 2017). Our current understanding of collective behaviour is however incomplete from a developmental perspective because there is limited information on the behavioural changes that occur in animal collectives over short timescales without modification to environmental conditions (Bengston and Jandt, 2014; Biro et al., 2016; Hinz and de Polavieja, 2017; Sasaki and Biro, 2017). Increased attraction between individuals during early development is one such mechanism by which this can occur (Hinz and de Polavieja, 2017; Masuda and Tsukamoto, 1998). Alternatively, changes in collective behaviour can be associated with repeated exposure to the same environmental conditions, such as acclimatisation to an environment that was initially novel. For example, zebrafish shoals spend an increasing proportion of time in a disordered compared to an ordered state as they habituate over time to a novel environment (Miller and Gerlai, 2012).

Different groups within the same population can also consistently differ in their collective behaviour over time and contexts. This group-level variation is akin to the consistent behavioural variation between individuals described as animal personality variation (Réale et al., 2007). Evidence for consistent differences between groups, often referred to as group or collective personality variation, is strongest in social insects in behaviours associated with group defence and foraging (e.g. Pinter-Wollman, 2012; Planas-Sitja et al., 2015), and has been linked to colony productivity and survival (Wray et al., 2011). There are also examples in vertebrates, for instance, in both chimpanzees (van Leewen et al. 2018) and fish (Jolles et al. 2018).

Not only do individuals vary in their average levels of behaviour, but they may also vary in their change in behaviour over time or contexts; i.e. they vary in their behavioural plasticity. Individual red knots, for example, differ in their change in vigilance behaviour in response to predation risk (Mathot et al., 2011). Differences in plasticity can be quantified using a behavioural reaction norm approach where random slope parameters in regression models are free to vary among individuals or groups (Dingemanse et al., 2010; Nussey et al., 2007; van de Pol and Wright, 2009). While reaction norm approaches have been applied widely to study variation in plasticity between individuals, they have not been used to examine differences between groups in the plasticity of collective behaviour. Examination of random slope variation is, however, key to understanding the emergence and maintenance of consistent group-level differences in behaviour because differences between groups may be lost or magnified over time.

It thus remains unclear when variation between groups is apparent; inter-group differences could establish early in group formation, and be stable or diminish over time, or they could develop gradually. In three-spined sticklebacks, fish form shoals of unrelated individuals (Peuhkuri and Seppä, 1998) and group membership may remain relatively stable over time (Ward et al., 2002). Here, we tested twelve groups of eight sticklebacks repeatedly in the same context (an open arena) under controlled laboratory conditions. The groups were tested over multiple days up to twelve times over four weeks. Each trial lasted approximately 25 minutes and group behaviour was analysed during five 2.5-minute time intervals that were separated by the presentation of a food item (see Materials and Methods, and MacGregor et al., 2020 for analyses of responses to the food item presentations). We used high resolution video tracking to examine changes over time in six traits of collective motion that are known to be functionally important: polarisation (where higher levels of polarisation can improve social information transfer and reduce the risk of predation due to confusion effects: Attanasi et al., 2014; Ioannou et al., 2012), the speed of the group centroid (where group speed can affect responses to predators and encounter rates with food: Wood and Ackland, 2007), convex hull area as a measure of spacing between individuals (where less cohesive groups are at greater risk of predation: Duffield and Ioannou, 2017), the rate of transitions in state between polarised and unpolarised swarm-like states (Calovi et al. 2015; Tunstrøm et al., 2013), the maximum cross-correlation in speed (which is a measure of information transfer: Katz et al., 2011), and the rate of switches in leadership over time (where individuals at the front of the group have greater access to foraging rewards at a cost of greater risk of predation, and increased switching is observed in shoals from high predation habitats: Burns et al., 2012; Herbert-Read et al., 2019; Ioannou et al., 2019; Reebs, 2000). Using a reaction norm approach combined with model selection procedures (Dingemanse et al., 2010; Symonds and Moussalli, 2011), we then determined whether the groups differed from one another in the dynamics of their collective behaviour over shorter (within trials) and longer (between days) timescales.

## Results

To test for an effect of time on collective motion, each movement trait was analysed as a response variable in a separate linear mixed model (LMM), except for the rate of switches in leadership, which was analysed in a generalised linear mixed model (GLMM) fitted with a beta distribution. As time progressed both within trials (increases in interval) and between days, the groups became slower, less polarised and less cohesive on average. Individuals also became less responsive to their near neighbours, with lower maximum cross-correlations in speed (Table 1). Although the groups transitioned between polarised and unpolarised states significantly more at the end compared to the start of the trials, this effect was not observed between days (Table 1). There was no evidence for overall change in switches in leadership over time either within trials or between days (Table 1).

**Table 1.** Results from statistical tests of the main effects of interval within trial and between days on each trait. Traits were analysed separately in (G)LMMs with group identity included as a random intercept. Convex hull area was square root transformed to meet parametric assumptions. Model parameter estimates ± standard error (SE) are reported for each predictor.

|  | Interval |  |  | Day |  |  |
| --- | --- | --- | --- | --- | --- | --- |
| Trait | Estimate ± SE | Test Statistic | P | Estimate ± SE | Test Statistic | P |
| Polarisation | -0.023 ± 0.0034 | $F_{1,402} = 45.9$ | < 0.0001 | -0.012 ± 0.019 | $F_{1,412} = 39.0$ | < 0.0001 |
| Centroid speed | -0.41 ± 0.038 | $F_{1,402} = 116$ | < 0.0001 | -0.13 ± 0.0021 | $F_{1,410} = 37.1$ | < 0.0001 |
| Convex hull area | 19 ± 3.08 | $F_{1,402} = 38.5$ | < 0.0001 | 15 ± 1.73 | $F_{1,411} = 79.1$ | < 0.0001 |
| Transitions in state | 0.0026 ± 0.00023 | $F_{1,402} = 127$ | < 0.0001 | -0.000084 ± 0.00013 | $F_{1,412} = 0.41$ | 0.52 |
| Maximum cross-correlation in speed | -0.0085 ± 0.0020 | $F_{1,402} = 17.8$ | < 0.0001 | -0.014 ± 0.0011 | $F_{1,412} = 145$ | < 0.0001 |
| Switches in leadership | -0.00011 ± 0.0094 | $\chi^2_1 = 0$ | 0.99 | -0.009 ± 0.0051 | $\chi^2_1 = 3.3$ | 0.069 |

To determine whether there were consistent differences in the average value of each trait between groups, we compared the performance of models fitted with and without group identity as a random intercept (Table 2). Based on a difference in Akaike Information Criterion of greater than two units (Symonds and Moussalli, 2011; Vaida and Blanchard, 2005), models including group identity were more likely given the data than models without for all traits, suggesting that the groups differed in their collective motion (Table 2).

**Table 2.** Conditional AIC (cAIC) and corrected AIC (AICc) scores for models with different random effect structures (with and without group identity as a random intercept and with and without interval and day as random slopes). All models included interval and day as main effects. Estimated degrees of freedom (DF), Δ(c)AIC(c) from the (c)AIC(c) of the best supported model (in bold) and model rank (where 1 is the best supported and 4 the least supported model) are reported for each trait. AICc is estimated for models with no random effects and used in the comparison of models of the rate of switches in leadership because cAIC is unavailable for beta regressions. Convex hull area was square root transformed to meet parametric assumptions. Missing values are where models were unable to estimate intercept-slope correlation parameters and therefore produced unreliable (c)AIC(c) scores.

| Trait | Random Intercept | Random Slope | (c)AIC(c) | DF | $\Delta(c)AIC(c)$ | Rank |
| --- | --- | --- | --- | --- | --- | --- |
| Polarisation | None | None | -598.29 | 4 | 267.25 | 4 |
|  | Group Identity | None | -731.37 | 14.4 | 134.17 | 2 |
|  | Group Identity | Interval | -726.29 | 17.6 | 139.25 | 3 |
|  | **Group Identity** | **Day** | **-865.54** | **24.0** | **0** | **1** |
| Centroid speed | None | None | 1438.30 | 4 | 321.21 | 4 |
|  | Group Identity | None | 1271.81 | 14.6 | 154.72 | 2 |
|  | Group Identity | Interval | 1274.47 | 19.7 | 157.38 | 3 |
|  | **Group Identity** | **Day** | **1117.09** | **24.5** | **0** | **1** |
| Convex hull area | None | None | 5075.68 | 4 | 323.27 | 3 |
|  | Group Identity | None | 4927.27 | 14.5 | 174.86 | 2 |
|  | Group Identity | Interval | - | - | - | - |
|  | **Group Identity** | **Day** | **4752.41** | **24.7** | **0** | **1** |
| Transitions in state | None | None | -2848.1 | 4 | 124.70 | 3 |
|  | Group Identity | None | -2968.5 | 14.3 | 4.28 | 2 |
|  | **Group Identity** | **Interval** | **-2972.8** | **20.6** | **0** | **1** |
|  | Group Identity | Day | -2968.5 | 19.3 | 4.28 | 2 |
| Maximum cross-correlation in speed | None | None | -1056.5 | 4 | 226.88 | 4 |
|  | Group Identity | None | -1177.3 | 14.3 | 106.16 | 2 |
|  | Group Identity | Interval | -1171.6 | 17.5 | 111.87 | 3 |
|  | **Group Identity** | **Day** | **-1283.4** | **23.6** | **0** | **1** |
| Switches in leadership | None | None | -5085.3 | 4 | 16.72 | 2 |
|  | **Group Identity** | **None** | **-5102** | **5** | **0** | **1** |
|  | Group Identity | Interval | - | - | - | - |
|  | Group Identity | Day | - | - | - | - |

There were also differences between the groups in how their collective motion changed over time (Figure 1, A-E). Comparing the performance of models with and without day and interval as random slope parameters, the most likely models explaining variation in polarisation, centroid speed, convex hull area and maximum cross-correlation in speed included day as a random slope (Table 2, Figure 1, A-D).

**Figure 1.**
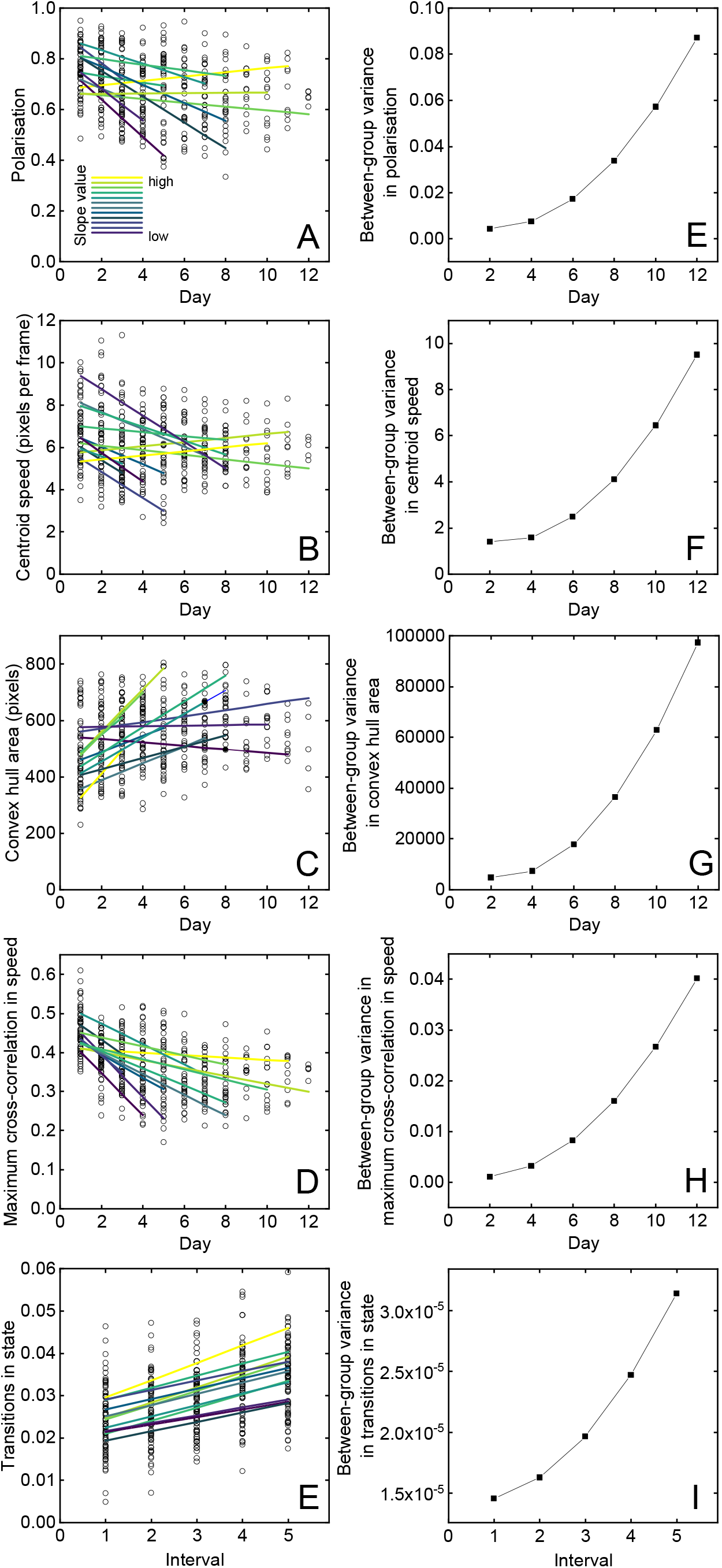
Variation between the groups in collective motion over time. Panels **A-D** show the predicted reaction norms of the twelve groups across days from the best supported models to explain polarisation, centroid speed, convex hull area and maximum cross-correlation in speed, and panel **E** shows the predicted reaction norms from intervals during a trial from the best supported model to explain the rate of transitions in state. The predicted reaction norms for each group are coloured by slope (highest (yellow) to lowest (dark purple)) within each panel and estimated using the *predict* function in R package *lme4* (Bates et al., 2015). For predictions, the interval (in panels **A-D**) and day (in panel **E**) are held at their mean value. The open circles are the raw values (i.e., per interval per trial, n = 416). For convex hull area, values are square root transformed to meet parametric assumptions. In **A-D** the lines are truncated to the day where the experiment ended for a particular group. Panels **F-H** show the trends in between-group variance over days of the experiment. Between-group variance is the model-estimated variance in the random intercept of group identity when the variable is centred at day two, four, six, eight, ten and twelve, respectively, shown as filled squares. Panel **I** shows the trend in between-group variance in the rate of transitions in state within trials for the model-estimated variance in the random intercept of group identity when the variable interval is centred at interval one to five, respectively. The trend in the model estimated variance in group identity reveals how inter-group differences in collective motion change over time.

While most groups showed less social collective behaviour for these traits as the days progressed (decreased polarisation, group speed and speed correlation, with an increase in convex hull area), some showed no change (Figure 1, A-D). As a result, the model estimated variance between the groups also increased over the days of the experiment, i.e. the differences between groups became more pronounced (Figure 1, F-I). Groups did not differ in how they changed over time within trials in these traits (Table 2). In contrast, the best supported model for the rate of transitions in state included interval as a random slope, showing that the groups differed in their response over a short timescale (Figure 1, E) and the model estimated variance between the groups also increased over time within trials (Figure 1, J).

## Discussion

Differences between groups in their collective behaviour may arise due to differences between individuals that are present when groups first form and then remain stable over time. Alternatively, variation between groups may become stronger as individuals in newly formed groups have opportunities to gather information and socially interact (Bengston and Jandt, 2014). Our data support the latter scenario with more variation between the groups in collective motion at the end compared to the start of the experiment. The groups displayed different rates of change in their collective motion, i.e. differing degrees of plasticity, with variation in the strength, and in some cases direction, of their response as the days of the experiment progressed. Previous work has reported consistent group differences across contexts in alignment, speed and cohesiveness in three-spined sticklebacks (Jolles et al., 2018 although see Ginnaw et al., 2020). We show that differences in these traits, as well as the maximum cross-correlation in speed, between shoals is magnified over several weeks when group membership is stable. Positive feedback through repeated interactions between the same individuals and learning can exaggerate initially small differences over time (Bengston and Jandt, 2014; Biro et al., 2016). This may explain how differences between groups in their collective behaviour over time may emerge from minimal between-group heterogeneity, which we ensured with the method used to assemble the groups.

Despite these differences between groups in their responses over time, overall, the groups tended to become slower, less cohesive and less polarised with reduced cross-correlation in speed both within trials and between days of testing. This is consistent with a weakening of attraction and responsiveness to near neighbours. Functionally, these temporal changes are associated with a reduction in social information sensing and anti-predator benefits (Ioannou et al., 2012; MacGregor et al., 2020; Partridge, 1982), and suggest that the groups were acclimatising over time to the low risk, predator free context. Maintaining social responsiveness during collective motion by forming fast, cohesive and highly polarised groups may be cognitively demanding and energetically costly (Di Santo et al., 2017). Therefore, in the absence of environmental pressures to preserve these traits of shoaling, animals become less collective in their movement (Miller and Gerlai, 2012).

Studies of animal collective motion typically only provide a snapshot of group behaviour. Demonstrating that groups show variation in their plasticity over time has significant implications for the study of collective behaviour as it magnifies variation between groups, making the detection of differences between treatments (Ginnaw et al., 2020), populations (for example along an environmental gradient; Herbert-Read et al., 2019) or species (Partridge et al., 1980) more difficult. A greater number of independent experimental groups may be required than originally thought, and reliance on a few replicate groups may be inadequate. Testing newly assembled groups may alleviate this, although this may not maximise ecological relevance if the animals have relatively stable social memberships. The time available for behaviour between groups to diverge will be determined by the degree to which social membership is stable, which in turn may be driven by ecological factors such as population density (determining the rate at which groups encounter one another and can exchange group members: Beauchamp, 2011) and predation risk (determining the likelihood individuals change groups: Hasenjager and Dugatkin, 2017; Heathcote et al., 2017). A more holistic understanding of collective motion would consider such ecological factors, as well as the development of collective behaviour over time.

## Materials and methods

### Ethics

All procedures regarding the use of animals followed United Kingdom guidelines and were approved by the University of Bristol ethics committee (UIN UB/17/060).

### Study animals

Three-spined sticklebacks (*Gasterosteus aculeatus*, 27 ± 2.4 mm, mean ± sd, standard body length at the time of testing for n = 96 individuals), were collected from the river Cary, Somerset, UK (grid ref: ST 469 303) in September 2016 and transported to fish facilities at the University of Bristol, UK. The fish were housed in three glass tanks (70 cm (L) × 45 cm (W) × 37.5 cm (H)) of approximately 50 individuals for 10 months before testing and were fed daily with brine shrimp or defrosted frozen bloodworm (*Chironomid* sp. larvae). Photoperiod was a 11:13 h light:dark cycle and ambient temperature was maintained at 16°C to prevent the fish from entering reproductive condition (fish were not sexed). One week prior to the experiment we assigned the fish to twelve groups of eight inviduals in a randomised complete block design by repeatedly netting twelve individuals from a tank and randomly allocating each individual to one of twelve groups. This minimised any potential differences between groups in case the netting of individuals from the holding tanks was biased towards sampling individuals with particular traits (Carter et al., 2012). The fish were given six days to habituate in their groups in smaller glass holding tanks (70 (L) × 25 (W) × 37.5 (H) cm) enriched with a horizontal piece of PVC tubing and an artificial plant prior to the first trial.

### Experimental procedure

The groups were tested repeatedly up to twelve times over four consecutive weeks. Each group was tested every other day from Monday to Friday. Six groups were tested per day, therefore at the start of each week the twelve groups were randomly allocated to two sets of six groups and the set that would have trials on Monday, Wednesday and Friday was determined at random. Trials took place between 09:45 and 16:00 each day and the order of testing of the groups within a day was randomised. Each trial was conducted in a white oval-shaped open experimental arena (133.5 (L) × 72 (W) × 62 (wall height) cm) with a water depth of 10 cm which was maintained at the same temperature as the holding tanks. Because individuals may seek refuge in heterogenous environments, no artifitial enrichment was provided in the arena to encourage collective movement. There was a small amount of variation in the length of each trial (25 ± 2.2, mean ± sd trial length in minutes for 84 trials) due to the experimental protocol (see MacGregor et al. 2020 for further details). Trials were filmed from above with a Panasonic HC-VX980 video camera in 4K (3840 × 2178 pixels) and a temporal resolution of 25 frames per second. After each trial the fish were returned to their holding tank and were fed with bloodworm following the final trial of each day. Trials were terminated for a group before the end of the fourth week if any individual in the group began displaying signs of poor health.

At the start of a trial, all eight fish were netted into the centre of the arena and allowed 2 minutes to acclimatise. For the part of the experiment analysed and reported in MacGregor et al. (2020), a red-tipped pipette that delivered a single food item (a defrosted bloodworm) was presented to each group on six occasions during each trial. The time interval between presentations was approximately 4 minutes, which allowed the group to resume normal swimming behaviour. In the present study, we analysed the video footage of the groups’ behaviour in the 2.5 minutes immediately prior to the second to sixth presentations of the red-tipped pippette in each trial to obtain trajectory data for five consecutive time intervals per trial (i.e. 12.5 minutes of video footage per trial). Data for three intervals from two separate trials were not obtained due to corrupted video files. During the first week of trials, one individual in each of three groups was replaced (one due to injury and two deaths of unknown cause) with an individual that was naive to the experiment and given 24 hours to habituate within their group in the holding tank prior to testing. All trials conducted prior to these replacements were included in the analyses.

### Data processing

Video files were converted to MPEG-4 HD (1920 × 1080 pixels) in Handbrake (version 1.0.7, https://handbrake.fr/) where 1 mm was equal to 2.7 pixels. Trajectory data for each fish were obtained using idTracker version 2.1 (Perez-Escudero et al., 2014). Only frames with complete trajectory information for all eight individuals in the groups were included in analyses. Frames where the speed of any individual in the group exceeded 50 pixels per frame (~ 46 cm s^−1^) were excluded from further analyses because visual inspection of the data revealed that these were likely due to tracking errors. The remaining trajectory data were then smoothed using a Savitzky-Golay filter with a span of 0.5 s (~13 frames) and a polynomial of 3 degrees in R package *Trajr* (McLean and Skowron Volponi, 2018). One interval in a trial had < 30 s of trajectory information after data processing due to poor tracking quality and was excluded. The final dataset consisted of trajectory information for 416 intervals (3705 ± 173, mean ± sd frames per interval) from 84 trials across the twelve groups.

We calculated the median (across frames for each interval per trial) of the groups’ polarisation, centroid speed based on the distance travelled since the previous frame, and the convex hull area due to the right skew in the distributions of these variables. For each interval, we also calculated the maximum cross-correlation in speed of an individual to their nearest neighbour (with a maximum lag of 300 frames (12 seconds)) and used the mean maximum cross-correlation in speed across the eight individuals for analyses. The rate of transitions in state was quantified by the number of transitions of the group between polarised (> 0.65) and unpolarised (< 0.65) states divided by the total number of frames in the time interval (excluding time steps with missing trajectory data). Finally, we quantified the rate of switches in leadership as the number of switches of each fish to or from the lead fish position in the group (determined by an individual’s rank distance along the direction of group motion (1st to 8th)) across consecutive frames when the group was in a polarised (> 0.65) state divided by the number of frames and used the mean rate of switches in leadership across the eight individuals per interval per trial for analyses. We restricted assignment of shoal position and hence calculation of switches in leadership to only those frames where the group was polarised (> 0.65) because here the groups had a clear direction of travel. Note that if a pair of fish switched their positions, this was counted as two switches, one for each fish.

The correlations between the traits are reported in Table S1. Because increases in convex hull area may reflect either the splitting of the group into subgroups or increases in inter-individual spacing between group members within a single group, we also report the correlation between the median convex hull area per interval and the median mean nearest neighbour distance per interval where the mean nearest neighbour distance is calculated across the eight individuals per frame. The high correlation between these variables suggests that convex hull area reflects the latter scenario.

### Data analyses

All data analyses were carried out in R version 3.6.2 (R Core Team, 2019). To test for an overall effect of time on collective motion, each model included interval (1st to 5th) and day (1st to 12th) as continuous main effects and group identity as a random intercept. Convex hull area was square root transformed to meet parametric model assumptions. Gaussian models were fitted using restricted maximum likelihood and the significance of interval and day were tested using Kenward-Roger approximation F-tests in R package *lmerTest* (Luke, 2016). The significance of the effects of interval and day on the rate of switches in leadership were tested with likelihood ratio tests with models fitted using maximum likelihood. Residuals from all models were checked visually for conformity to assumptions of homogeneity of variance and normality of error.

To determine whether the groups differed on average in their collective motion, the performance of models fitted with and without group identity as a random intercept (which accounts for average group-level variation) were compared using the cAIC (Vaida and Blanchard, 2005) and AICc, respectively. When the model with group identity was greater than two Δ(c)AIC(c) units from the model without, this was deemed as support for inter-group differences in behaviour (Symonds and Moussalli, 2011). cAIC controls for the number of effective model parameters accounting for shrinkage in the random effects (Vaida and Blanchard, 2005) and was estimated in R package *cAIC4* (Safken et al., 2018). For models without group identity (i.e. no random effects), the AICc of the model estimated in R package *AICcmodavg* (Mazerolle, 2016) was used for comparison. For switches in leadership models were compared based on AICc because cAIC estimates for beta regression models were unavailable.

Random slopes are interactions between fixed effect covariates and random effects, and represent reactions of each groups’ collective behaviour over time (Dingemanse and Dochtermann, 2013). To test whether the groups differed in their trajectories of change over time, the performance of models fitted with and without interval and day as random slopes (which account for group-level behavioural plasticity within trials and between days, respectively) were compared using the differences in cAIC. Three models (Table 2) were not included in model comparisons because (c)AIC(c) estimates were unreliable due to model overfitting that was indicated by estimated perfect correlations between random effect terms (Harrison et al. 2018). All models were fitted using maximum likelihood. Random slope variation disrupts repeatability which will vary depending on the level of the covariate, therefore we do not report inter-group repeatability estimates.

## Acknowledgements

This research was funded by a Natural Environment Research Council grant no. NE/P012639/1 awarded to CCI.

## Competing Interests

The authors declare no competing interests.

---

**Table S1.**
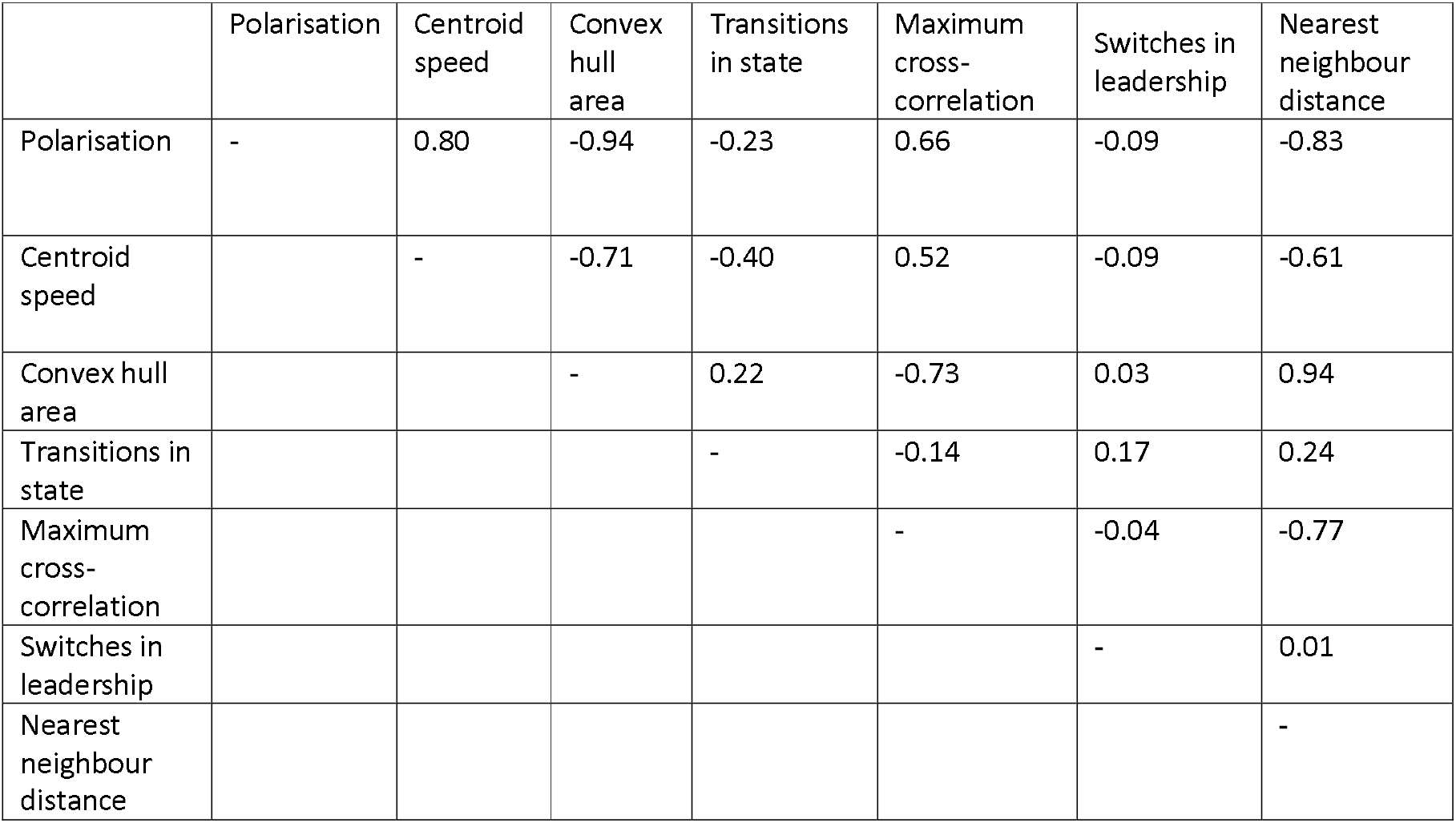
Spearman’s rank correlations between the six traits and median mean nearest neighbour distance.

|  | Polarisation | Centroid speed | Convex hull area | Transitions in state | Maximum cross-correlation | Switches in leadership | Nearest neighbour distance |
| --- | --- | --- | --- | --- | --- | --- | --- |
| Polarisation | - | 0.80 | -0.94 | -0.23 | 0.66 | -0.09 | -0.83 |
| Centroid speed |  | - | -0.71 | -0.40 | 0.52 | -0.09 | -0.61 |
| Convex hull area |  |  | - | 0.22 | -0.73 | 0.03 | 0.94 |
| Transitions in state |  |  |  | - | -0.14 | 0.17 | 0.24 |
| Maximum cross-correlation |  |  |  |  | - | -0.04 | -0.77 |
| Switches in leadership |  |  |  |  |  | - | 0.01 |
| Nearest neighbour distance |  |  |  |  |  |  | - |

## Notes

### Competing Interest Statement

The authors have declared no competing interest.

